# Metabolites from plasma-like medium fuel nitrogen metabolism and influence proliferation in *Leptospira interrogans*

**DOI:** 10.64898/2026.03.12.711193

**Authors:** Matthew H. Ward, Nathan Scherer, Leah P. Shriver, Gary J. Patti

## Abstract

Leptospirosis, caused by pathogenic *Leptospira* spp. such as *L. interrogans*, is a bacterial zoonosis of increasing prevalence with no consistently effective treatments in severe cases. We sought to characterize metabolic mechanisms that support *L. interrogans* infection in the host setting, with the ultimate goal of revealing unexplored therapeutic opportunities. We first established and validated a culture medium, which we refer to as supplemented Human Plasma-Like Medium (sHPLM). sHPLM more closely resembles the physiological environment of the human host than standard culture media, such as the EMJH (Ellinghausen-McCullough-Johnson-Harris) medium typically used for *Leptospira* culture. To better understand bacterial metabolism, we pioneered metabolomics in sHPLM-cultured *Leptospira*. Specifically, we developed a liquid chromatography mass spectrometry (LC/MS) metabolomics-based workflow for both medium analysis and stable isotope tracing with *L. interrogans* cultures. The application of these innovations revealed that the amino acid glutamine is a major nitrogen source for *L. interrogans*. A small-molecule inhibitor blocking glutamine utilization, JHU-083, effectively impaired the proliferation of sHPLM cultures. Further, adding glutamine to non-physiological EMJH medium rapidly induced a short-term proliferative boost in *L. interrogans* and increased biofilm formation. RNA-sequencing after glutamine exposure revealed transcriptional trends for increases in biosynthesis to support these phenotypes. Although ammonium has long been thought to be the sole nitrogen source for *L. interrogans,* our results demonstrate that glutamine provides a second source of nitrogen for biosynthesis and may act as a metabolite signal to alter *L. interrogans* physiology in ways that could influence infection. This work highlights that studying *L. interrogans* under physiological conditions is key to understanding mechanisms supporting infection and points to nitrogen assimilation as a potential target for therapies.

**Author Summary:** Leptospirosis is a potentially fatal disease transmitted through water and soil contaminated with pathogenic *Leptospira* bacteria. Much research is currently focused on the idea that an improved understanding of how *Leptospira* infects hosts and causes disease may inspire the development of improved therapeutics, which are urgently needed. Focusing on *Leptospira interrogans*, a clinically important pathogenic species, we determined that conventional growth media are inadequate for understanding how the bacterium behaves when inside hosts. Instead, we designed an optimized formulation to mimic human blood, and we applied an underutilized technique for measuring the biochemical reactions that enable pathogen survival. These two innovations revealed that *L. interrogans* uses glutamine, an abundant nutrient in host blood and tissues, as a source of nitrogen for the production of biomolecules that are required for replication and infection. This discovery is notable as nitrogen demands were previously thought to be met using ammonium. Treating *L. interrogans* with inhibitors of both glutamine and ammonium metabolism blocked bacterial replication. We also discovered that *L. interrogans* increases its growth rate, upregulates its expression of biosynthetic pathways when exposed to glutamine, and increases its formation of biofilm. Our results reveal the importance of glutamine in supporting the lifecycle of leptospirosis-causing bacteria.

## Introduction

Leptospirosis is a bacterial disease associated with exposure to flooding and poor sanitation^1,2^. Incidence worldwide reached an estimated annual count of one million in 2015 and continues to steadily increase across many regions, with outbreak fatality rates between 5-20%^3–5^. Preclinical interventions conducted post-colonization and meta-analyses of clinical studies have called into question the effectiveness of standard-of-care antibiotics, including penicillin, outside of prophylaxis^6,7^. Some of these antibiotics target a specific area of cellular metabolism (e.g., peptidoglycan biosynthesis); however, there are many other pathways that are also critical for replication, motility, host immune evasion, and toxin production^8–11^ that have not been evaluated for therapeutic viability in leptospirosis. Metabolic interventions outside of cell wall biosynthesis have been successful in the treatment of other infectious diseases, such as tuberculosis^12^. We posit that a better understanding of the metabolic mechanisms supporting infection and pathogenicity of disease-causing *Leptospira* spp. could reveal new, urgently needed treatment strategies.

Early work demonstrated that pathogenic *Leptospira* spp., such as *Leptospira interrogans*, require exogenous fatty acids as a carbon source for biomass and energy production^13^. Similar studies revealed that carbon from fatty acids and select amino acids is used by pathogenic *Leptospira* spp. to biosynthesize proteins as well as pyrimidine and purine nucleotides^14^. In contrast, the comparative preferences for different nitrogen sources in *Leptospira* spp. are unknown. The specific nitrogen assimilation reactions and the molecules into which nitrogen is incorporated also remain unevaluated. Instead, past studies concluded that ammonium was the preferred source of nitrogen based on growth rate^14,15^. Given that protein-based toxins, nucleic acids, and polyamines all contain nitrogen and contribute to pathology and proliferation, nitrogen metabolism is likely a key driver of virulence and a potential vulnerability.

Liquid chromatography mass spectrometry (LC/MS)-based metabolomics enables mechanistic investigation of metabolism through pathway intermediate and activity measurements, but it has yet to be implemented in *Leptospira* research. Much of our modern understanding of the biochemical activities in *Leptospira* spp. has been inferred from genomics^16^, transcriptomics^17,18^, or proteomics^19,20^ data. Accordingly, this body of knowledge needs to be confirmed with direct measurements. Moreover, most metabolism studies in *Leptospira* spp. have used non-physiological media that do not replicate the nutrient, temperature, or osmolarity conditions of human hosts. As a result, and bolstered by reports of changes in metabolic genes during host adaptation^17,18,21–23^, it is unclear whether past conclusions about metabolism translate accurately to *in vivo* settings.

Here, we developed approaches to overcome both of the above barriers that have hindered progress to understand and therapeutically exploit *Leptospira* spp. metabolism. Inspired by efforts in cancer research to create physiologically relevant media^24,25^ and building off of recent work in *Leptospira*^26^, we designed a medium that closely reflects the nutrient conditions experienced by *Leptospira* spp. in human hosts. In parallel, we also established LC/MS-based metabolomics and isotope-tracing workflows for *Leptospira* spp. The application of our updated medium formulation and global profiles of metabolic pathways led us to the surprising observation that glutamine (Gln) is a major source of nitrogen for *Leptospira* spp. and that it can control the rate of proliferation and biofilm formation.

## Results

### Establishing a workflow to study *L. interrogans* metabolism

Recent work in both mammalian cells and microbes has shown that the physiology of cells cultured in conventional medium formulations that have minimal components at supraphysiological concentrations does not reflect the physiology of the same cells *in vivo*^18,24,26^. Concerns over this observation led to the development of human plasma-like medium (HPLM, Gibco, A4899101) containing a greater variety of small molecules at concentrations similar to average values in human plasma^27,28^. While this formulation is not intended for long-term growth without medium changes, because nutrients may be rapidly depleted by highly active cell types, it improves confidence in the translatability and accuracy of research findings. Notably, lipophilic metabolites normally present in human plasma are not included in the current commercially available formulation.

We aimed to develop a physiologically relevant medium for *Leptospira* culture. Because *Leptospira* spp. require fatty acids for carbon assimilation^29^ and to improve the physiological relevance of HPLM, we first attempted continuous culture in HPLM supplemented with 10% fetal bovine serum (FBS) and a bovine serum albumin (BSA):oleate conjugate (**Fig 1A**). We inoculated the supplemented HPLM (sHPLM) with the commonly used strain Fiocruz L1-130 of *Leptospira interrogans* serovar Copenhageni, which causes hepatic, pulmonary, and renal pathology in hamsters^30^ and C3H/HeJ mice^31,32^. Conventionally, *Leptospira* strains are cultured at 28-31 °C in EMJH (Ellinghausen-McCullough-Johnson-Harris) medium, a serum-lacking minimal medium. Ammonium, supplied at 4.67 mM^33^, is the sole nitrogen source in EMJH, which is in 100-fold excess compared to most locations in host organisms from which *L. interrogans* can be isolated. Tween 80, a xenobiotic surfactant carrying a mix of fatty acids, primarily oleate, is the carbon source. In such non-physiological conditions, the growth rate is typically 18 hours per doubling^7^. Experimentally with EMJH, we observed an average of 15.5 hours per doubling during exponential phase (**S1A Fig**). By contrast, in sHPLM cultures we observed robust growth, reaching a maximum rate of approximately 4.5 hours per doubling when cultured at 37 °C and 5% CO_2_. This growth rate is about twofold greater than values obtained *in vivo* using qPCR and luminescent *L. interrogans* in guinea pig and mouse models^7,34^, where host immune mechanisms would be opposing maximal replication. We also hypothesized that depletion of required nutrients would be growth-limiting as is expected for physiological media, and indeed by adding additional BSA:oleate conjugate we could increase the stationary phase density (**Fig 1A**).

**Figure 1.**
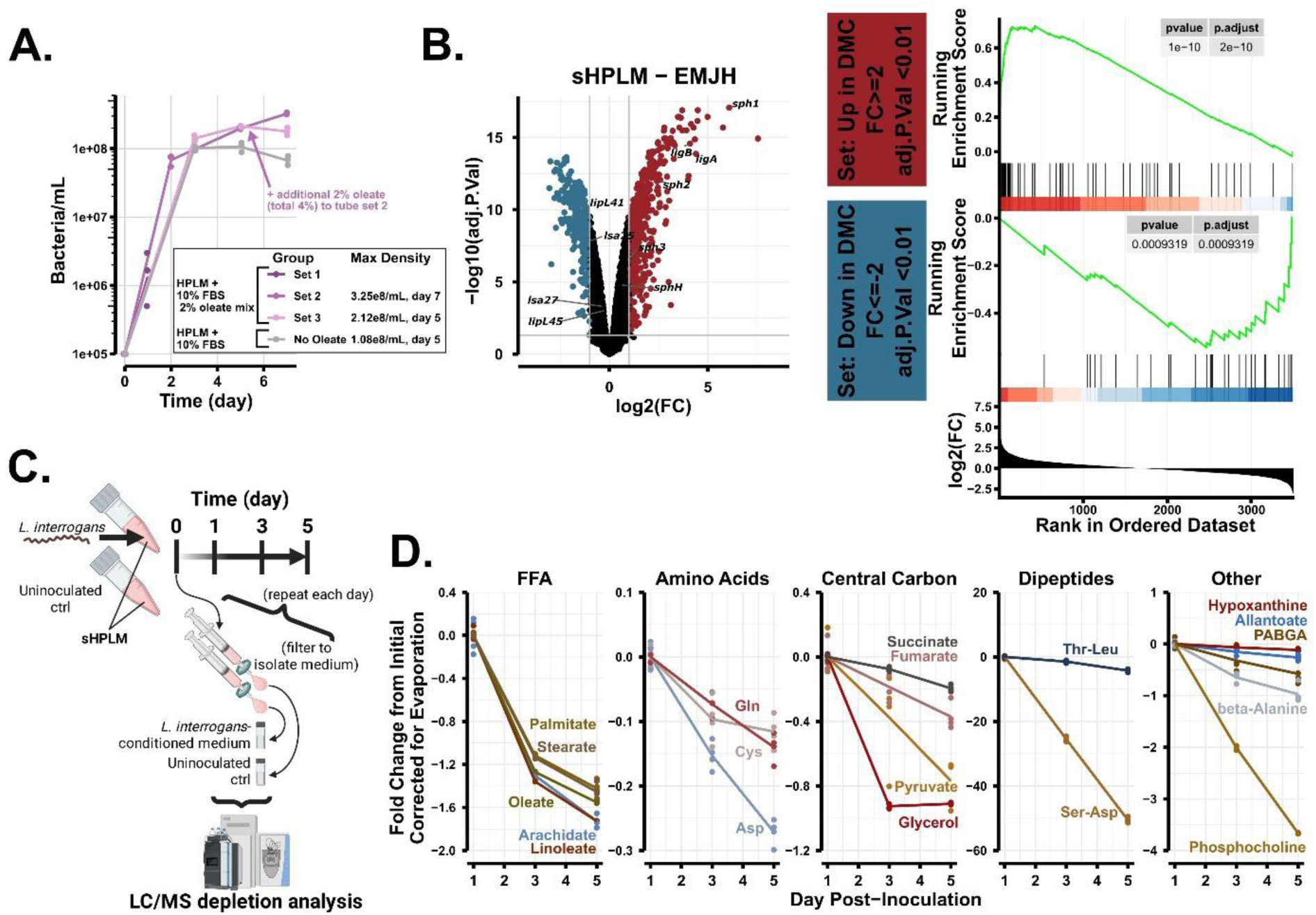
An LC/MS-based nutrient screen for *L. interrogans* using plasma-like medium. **A.** Darkfield-counted bacterial density over seven days in the indicated sHPLM mixture. Inoculation at 1e5 bacteria/mL in three sets of triplicate 5 mL tubes, indicated by color, with sets counted on different days to minimize effects of counting. **B.** RNA-sequencing differential expression analysis. (left) DEGs from *L. interrogans* exponential phase (5e7 bacteria/mL) EMJH cultures after four hours in EMJH or sHPLM. Red: increased transcript abundance; blue: decreased. Select virulence-related genes labeled. (right) Gene set enrichment analysis of fold change-ranked transcripts from the sHPLM-EMJH comparison using gene sets formed from false-discovery rate < 0.01 and −1≥ log2(FC) ≥1 DEGs in the Caimano et al. dataset^18^. **C.** Schematic for applying sHPLM and LC/MS to identify metabolites time-dependently depleted from host-like medium in *L. interrogans* monocultures. **D.** Metabolites with a fold change < −11% from days one to five. Sample fold change corrected against changes in uninoculated medium over time. n=3 independent samples per group for A, and n=4 for B and D. DEG, differentially expressed gene; DMC, dialysis membrane chamber; EMJH, Ellinghausen–McCullough–Johnson–Harris medium; FBS, fetal bovine serum; FC, fold change; FFA, free fatty acids; LC/MS, liquid chromatography mass spectrometry; PABGA, para-aminobenzoylglutamic acid; sHPLM, supplemented Human Plasma-Like Medium; Asp, aspartic acid; Cys, cystine; Gln, glutamine; Thr, threonine; Leu, leucine.

To further examine the extent to which our system modeled the host environment, we performed RNA-sequencing on *L. interrogans* four hours after switching to sHPLM (**Fig 1B** and **S1 Table**). Changes in the gene expression profile, compared to EMJH cultures, were similar to those observed in a published transcriptomics study evaluating *L. interrogans* isolated from living hosts against EMJH cultures^18^. We also observed specific gene changes in sHPLM-cultured *L. interrogans* consistent with host adaptation and virulence, including in *ligA*, *ligB*, *sph1/2/3*, *sphH*, *lsa27*, *lsa25*, *lipL41*, and *lipL45* (**S1B Fig**). Gene set enrichment and overrepresentation analysis revealed a trend for downregulation of biosynthetic metabolism, energy generation, and translation, as well as a trend for upregulation of protein import, fatty acid biosynthesis, and transcription (**S1C Fig** and **S1D**).

With a strategy in place for culture under host-like conditions, we next sought to explore the metabolic properties of *L. interrogans* by profiling which nutrients were utilized. We inoculated *L. interrogans* into sHPLM at mid-exponential phase and extracted metabolites from filtered conditioned medium after one, three, and five days of culture for LC/MS-based metabolomics (**Fig 1C**). We identified 111 metabolites in fresh and culture-conditioned sHPLM and measured the relative concentration of each as a function of time (**S2 Table**). Metabolites decreasing over time relative to uninoculated replicates were assumed to be nutrients taken up by bacteria. Accordingly, we report 19 putative nutrients that were depleted by *L. interrogans* when cultured under host-like conditions (adjusted p-value < 0.05 and fold change decrease greater than 11%). The fold change cutoff was defined by a split in the distribution of fold change values from day one to day five (**Fig 1D** and **S1E Fig**). Consistent with prior reports using EMJH^35^, we observed a depletion of several fatty acids and glycerol. Notably, we also identified the depletion of three amino acids: cysteine (Cys), aspartic acid (Asp), and glutamine (Gln). The observation that not every amino acid was significantly depleted (**S1F Fig**) suggests roles for Cys, Asp, and Gln outside of proteinogenesis which, if required for *L. interrogans* survival under host-like conditions, could potentially be exploited to design novel therapeutics.

### Glutamine and ammonium are major nitrogen sources for *L. interrogans*

Nitrogen utilization in *L. interrogans* has been studied far less than carbon metabolism, yet it is equally important to virulence mechanisms. Of the hits from our depletion screen containing nitrogen and with reported concentrations, only Gln was supplied at a sufficiently high concentration to support biomass production. To evaluate the contribution of Gln to the nitrogen pool of *L. interrogans* relative to other potential nitrogen sources such as ammonium and urea^36^, we turned to stable isotope tracing.

We spiked sHPLM with a given ^15^N-labeled nutrient (the tracer) at 50% (i.e., a 1:2 ratio) of the existing non-labeled nutrient concentration in HPLM, for a starting value of 33% labeling in the extracellular metabolite pool. This amount was expected to minimize the perturbation of metabolic steady state while still providing enough isotope label to trace. We collected intracellular and extracellular samples with a combined workflow for analysis of tracer assimilation (**Fig 2A**), which relied upon syringe and vacuum filtration (more details in Methods). Our approach separated intracellular and extracellular metabolites faster than conventional methods that rely on centrifugation, which is critical for accurate analysis of rapid metabolic processes. As expected, vacuum-filtered culture (intracellular) samples showed much higher amino acid abundances than uninoculated controls without bacteria (**Fig 2B** and **2C**). Thus, this method enabled separation of the intracellular and extracellular amino acids, even those with high medium concentrations that would be expected to contaminate the intracellular sample otherwise.

**Figure 2.**
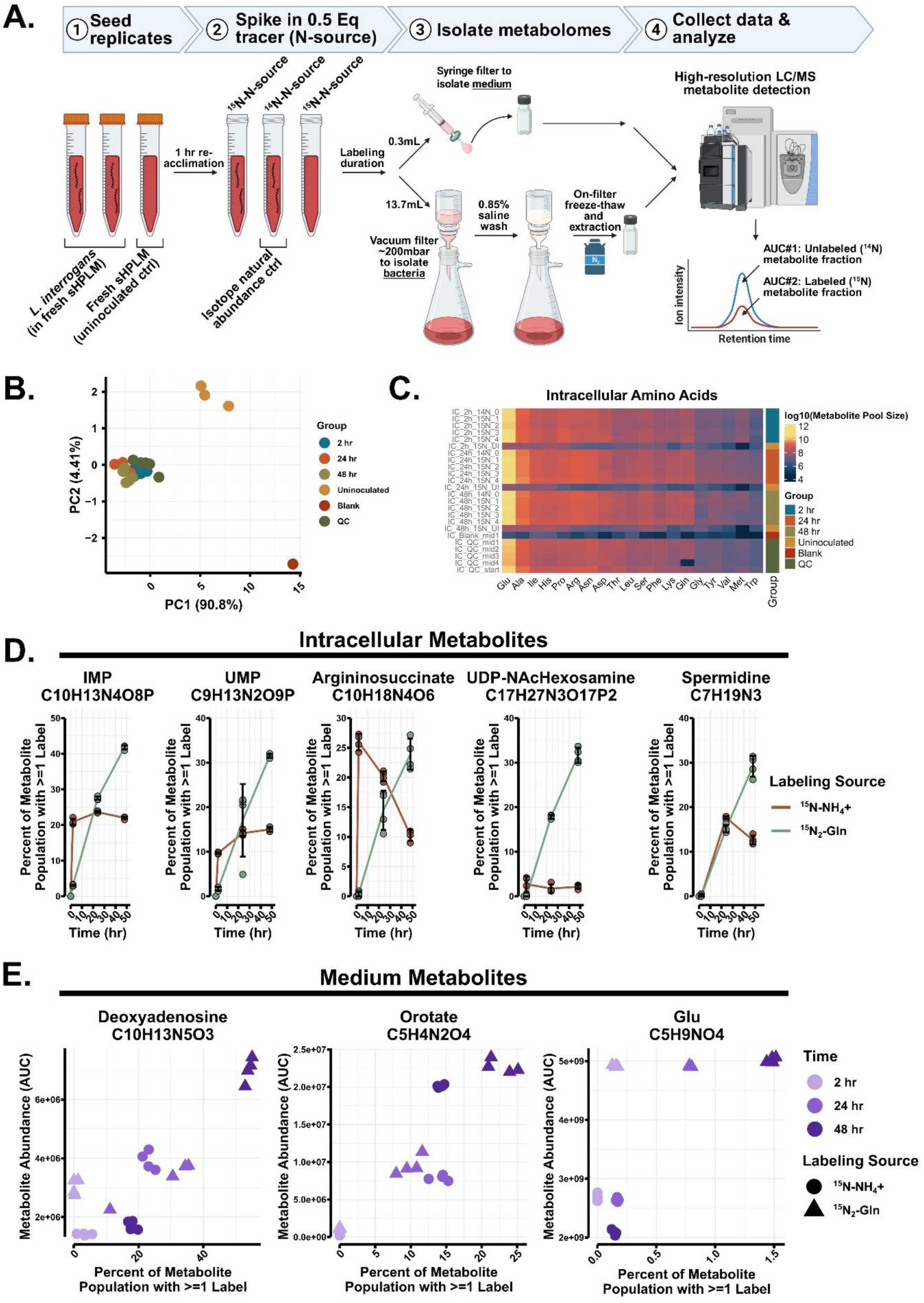
Ammonium and glutamine are *L. interrogans’* major nitrogen sources in physiological conditions. **A.** Schematic for combined stable isotope tracing analysis of intracellular and medium metabolomes. **B.** Principal component analysis plot and **C.** heatmap of summed pool sizes of 19 detected amino acids in samples processed through the vacuum filtration workflow. 2-, 24-, and 48-hour samples are intracellular *L. interrogans* metabolomes; uninoculated samples are time-matched filtered culture medium without bacteria; blank is extraction solvent only; QC is a pooled sample from all other samples. **D.** Percentage of given metabolite populations that have at least one labeled nitrogen from host-like medium-derived ammonium or Gln after 2, 24, or 48 hours of labeling in sHPLM spiked to a concentration of 33.3% labeled nitrogen source. Metabolites are representatives of the pathways: purine, pyrimidine, arginine, glycosylation, and polyamine biosynthesis. Zero-hour timepoint is imputed at 0% labeling. **E.** Media metabolite percentage labeling from the same experiments in D. n=4 independent samples per group for B-E. Ala, alanine; Arg, arginine; Asn, asparagine; Asp, aspartic acid; AUC, area under the curve; Cys, cysteine; Gln, glutamine; Glu, glutamic acid; Gly, glycine; His, histidine; Ile, isoleucine; IMP, inosine monophosphate; LC/MS, liquid chromatography mass spectrometry; Leu, leucine; Lys, lysine; Met, methionine; Phe, phenylalanine; Pro, proline; Ser, serine; sHPLM, supplemented Human Plasma-Like Medium; Thr, threonine; Trp, tryptophan; Tyr, tyrosine; Val, valine; UDP-NAcHexosamine, uridine diphosphate N-acetylhexosamine; UMP, uridine monophosphate.

After incubation with a ^15^N-labeled tracer, evidence of bacterial uptake and assimilation was assessed by the time-dependent accumulation of other biomolecules bearing ^15^N label at the nitrogen positions in their structures. These had been built through biochemical transformation of at least the nitrogen atoms from the tracer (**S2A Fig**). In time-resolved isotope tracing, the labeling percentage tends to increase until reaching a plateau. The parameters of this curve for a given biomolecule indicate the rate of tracer incorporation (flux), the size of the metabolite pool, and the contribution of the tracer to biosynthesis relative to other sources of nitrogen.

As a benchmark for comparison, we first cultured bacteria in sHPLM spiked with ^15^N-ammonium. Probable glutamine synthetase (GlnA) activity, the expected reaction assimilating ammonium, led to the labeling of predominantly one of two nitrogen atoms in glutamine (**S2B Fig**), and we also detected time-dependent increases of ^15^N labeling in downstream metabolites across several biosynthetic pathways (**Fig 2D** and **S3 Fig**). Our data provided evidence that host-derived ammonium fuels nucleotide biosynthesis (both purines and pyrimidines), amino acid biosynthesis, glycosylation, and polyamine biosynthesis in host-like conditions. Analysis of medium metabolites similarly showed time-dependently increased labeling of nucleic acids and amino acids, with some also increasing in total abundance as if secreted (**Fig 2E**). Consistent with the idea that these labeled metabolites in the medium were secreted during the experiment rather than leaked from bacterial lysis, other metabolites (e.g., hexose phosphates and NAD^+^) were only detected in intracellular samples (**S2C Fig**). These data suggest that select newly synthesized nucleic acids and amino acids are secreted by *L. interrogans*, potentially in the form of a short peptide or extracellular DNA.

We next compared these results against an identical experiment using ^15^N_2_-Gln as the tracer. Remarkably, we observed that in any given biomolecule population, the percentage that was built with Gln-derived nitrogen after one day of labeling was comparable or even higher than from ^15^N-ammonium in most observed areas of nitrogen metabolism (**Fig 2D** and **S3 Fig**). There was no pathway that exclusively utilized either nitrogen source, though some metabolites showed differential labeling dynamics. The lack of exclusivity in nitrogen flux could indicate metabolic redundancy. We note that although the synthesis of intracellular glutamine using nitrogen from exogenous ammonium generated primarily singly labeled Gln (M+1), little of this isotopologue was synthesized with nitrogen from exogenous ^15^N_2_-Gln (**S2B Fig**). This confirms that Gln is directly imported and incorporated into *L. interrogans* metabolism rather than first needing to be deaminated/deamidated to release ammonium. Similar to the ammonium labeling experiment, select metabolites in the medium were also labeled time-dependently with ^15^N_2_-Gln-derived nitrogen (**Fig 2E** and **S2C Fig**).

We performed similar isotope tracing experiments with ^15^N-Asp, due to its depletion from sHPLM, and ^15^N_2_-urea, a previously proposed nitrogen source^36^. In both cases, only minimal percentages of intracellular metabolites from sHPLM cultures were labeled (**S2D Fig**). These data suggest that Asp and urea are only minor sources of nitrogen under physiological conditions. We conclude that, contrary to conventional wisdom drawn from studies in non-physiological medium, *L. interrogans* does not rely solely upon ammonium for nitrogen assimilation. Instead, we show that Gln is an important source of nitrogen for *L. interrogans* when grown in host-like conditions with access to a variety of potential nitrogen sources. We also present a metabolic map of the major biosynthetic pathways active in *L. interrogans* under host-like conditions with data from which qualitative metabolic flux estimates can be drawn for future vulnerability analysis and the development of therapeutic interventions (**S3 Fig**).

### *L. interrogans* is sensitive to chemical inhibition of nitrogen import and metabolism

Our tracing studies yielded a model of parallel Gln and ammonium utilization for biosynthesis under host-like conditions. When grown in EMJH at 30 °C, both ammonium and Gln were sufficient as the sole nitrogen source to fuel proliferation at equivalent rates (**S4A Fig**), indicating that metabolic redundancy is possible. We next sought to discern whether ammonium and Gln were similarly redundant for proliferation in sHPLM. Simultaneously, because the literature and our RNA-sequencing indicated metabolic remodeling in *L. interrogans* grown in host conditions, we set out to reveal the mechanisms of nitrogen assimilation as well as host adaptation-associated changes by inhibiting key steps. For this purpose, we used small-molecule inhibitors, including the ammonium assimilation inhibitors dimethylammonium chloride (DMA) and methionine sulfoximine (MSO) as well as the Gln utilization inhibitor JHU-083 (**S4B Fig**).

We first tested DMA, a competitive inhibitor of the ammonium-specific permease AmtB. To validate ammonium assimilation inhibitors, we evaluated proliferation with and without inhibitor in *L. interrogans* that had been passaged in EMJH medium lacking the ammonium chloride component and complemented with either 2 mM ammonium chloride or L-glutamine. DMA, at ∼25 times the concentration of ammonium similar to reports in other organisms^37^, blunted proliferation in *L. interrogans* grown on ammonium but not on Gln (**Fig 3A**). A concentration of 2.5 mM DMA blocked proliferation to a similar extent in sHPLM (**Fig 3B**), where the ammonium concentration is ∼40 µM, which confirmed the assumption that DMA sensitivity is a function of medium ammonium concentration. Given evidence of metabolic redundancy between Gln and ammonium from our isotope tracing, this result suggested that ammonium was required for some critical non-metabolic process in host-like conditions specifically.

**Figure 3.**
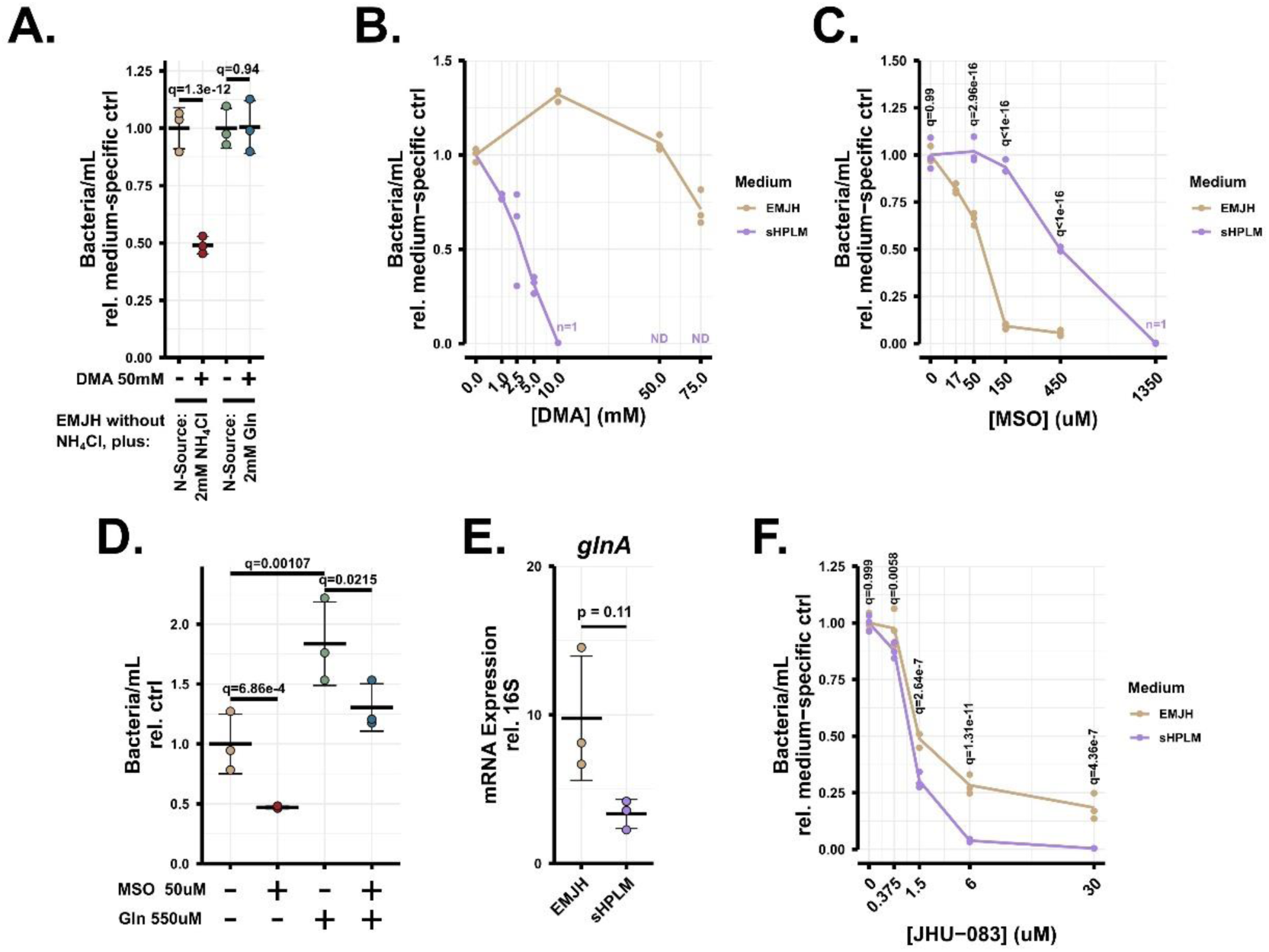
sHPLM cultures have differential sensitivity to glutamine and ammonium blockade. **A.** Proliferation assay of *L. interrogans* grown on custom EMJH (formulated without ammonium chloride) supplemented with either 2 mM ammonium chloride or Gln and treated with 50 mM dimethylammonium chloride to block ammonium transport competitively. Performed with darkfield microscopy counting after two days proliferating from 2.5e6 bacteria/mL seeding density. Replicates for each growth medium condition normalized to control for that growth medium. **B.** Dose-response two-day proliferation assay with DMA and *L. interrogans* in the indicated media, where ND indicated a density below the limit of detection (5e4 bacteria/mL). At 10 mM in sHPLM, only one replicate had detectable bacteria. X-axis in square-root scale. **C.** Proliferation assay measured at two days in the indicated culture medium, inoculated at 2.5e6 bacteria/mL with the indicated MSO concentrations. At 1350 µM in sHPLM, only one replicate had detectable bacteria. X-axis in square-root scale. **D.** Proliferation at two-day endpoint in standard EMJH (4.6 mM ammonium) supplemented with 50 µM MSO and/or 550 µM Gln. **E.** *glnA* mRNA levels in cultures passaged in EMJH or sHPLM for at least five days before RNA isolation. Expression relative to the housekeeping gene 16S rRNA of *L. interrogans*. **F.** Dose-response proliferation assay at two-day endpoint in response to varying JHU-083 dosages, inhibiting Gln-utilizing reactions. X-axis in square-root scale. n=3 independent samples per group for A-F. Welch’s t-test performed for E. A, C, D, and F multiple contrasts, see Statistics in methods. DMA, dimethylammonium; EMJH, Ellinghausen–McCullough–Johnson–Harris medium; Gln, glutamine; MSO, methionine sulfoximine; sHPLM, supplemented Human Plasma-Like Medium.

We next investigated the process of ammonium assimilation after import. The *L. interrogans* genome lacks an annotated homolog of glutamate dehydrogenase. If glutamate dehydrogenase activity is absent, the primary mechanism to assimilate ammonium nitrogen into biomass would be through synthesis of Gln from ammonium and glutamate by glutamine synthetase (GlnA) (**S4B Fig**). The glutamine synthetase inhibitor methionine sulfoximine (MSO) blunted proliferation in EMJH cultures of *L. interrogans* when grown on ammonium but not when grown on Gln (**S4C Fig**), suggesting that GlnA is indeed critical to EMJH cultures for the metabolic usage of ammonium. However, proliferation in sHPLM cultures was approximately nine-fold less sensitive to inhibition by MSO than standard EMJH cultures (**Fig 3C**). The most likely metabolite confounder to the effect of MSO, glutamic acid^38^, did not replicate the resistance of sHPLM cultures when added to EMJH cultures (**S4D Fig**). Further, supplementation with Gln and MSO together also did not block inhibition by MSO (**Fig 3D**), although Gln did independently increase endpoint density.

A more compelling explanation for MSO resistance in sHPLM was that metabolic remodeling unique to sHPLM-cultured *L. interrogans* led to upregulation of alternate pathways for ammonium assimilation (i.e., challenging the assumption of reliance on GlnA activity). To explore this line of reasoning, we examined the transcriptional expression of glutamine synthetase in sHPLM cultures. We observed weak evidence of *glnA* downregulation between EMJH and sHPLM cultures (**Fig 3E**). Querying our RNA-seq dataset from short-term sHPLM exposure revealed similar *glnA* downregulation, along with expression changes in associated nitrogen metabolism proteins and regulators (**S4E Fig**) solidifying a model of broad remodeling in ammonium assimilation mechanics. Synthesizing these results with the sensitivity to DMA, resistance to MSO, and rapid appearance of M+1 glutamic acid from ^15^N-ammonium (**S3 Fig**), we surmise that ammonium is metabolically but not completely redundant to Gln in sHPLM cultures and that changes in ammonium assimilation results in reliance on GlnA-independent mechanisms to fuel proliferation.

We last asked whether Gln incorporation was also necessary for sHPLM cultures. Notably, no Gln-specific permease has been described in *Leptospira* spp., such that Gln import is likely achieved through a multi-amino acid transporter. Therefore, we turned to the 6-diazo-5-oxo-L-nor-leucine (DON) prodrug JHU-083, which inhibits Gln-utilizing enzymes in mammalian cells^39^. We treated *L. interrogans* cultures with a range of JHU-083 concentrations and observed that proliferation was blunted by over half in the low micromolar range, with significantly higher sensitivity in sHPLM cultures (**Fig 3F**), once again signifying metabolic differences between sHPLM and EMJH cultures. We conclude that utilization of intracellular Gln for nitrogen donation is necessary for *L. interrogans* and could represent a therapeutic opportunity.

### Glutamine exposure induces a rapid proliferative bloom in *L. interrogans*

We observed that supplementing EMJH with Gln at physiological concentrations yielded *L. interrogans* cultures of higher density, implying that Gln increased their proliferation rate (**Fig 3D**). Before exploring the mechanism underlying this effect and its relation to the increased proliferation rate of sHPLM cultures, we examined whether it was conserved in two other *Leptospira* strains. We found that the addition of 550 µM Gln to EMJH cultures of *L. interrogans* serovar Manilae strain L495 did boost its proliferation at one day post-exposure. In contrast, the effect was not observed for the non-pathogenic, saprophytic *L. biflexa* serovar Patoc strain Patoc 1 (**Fig 4A**).

**Figure 4.**
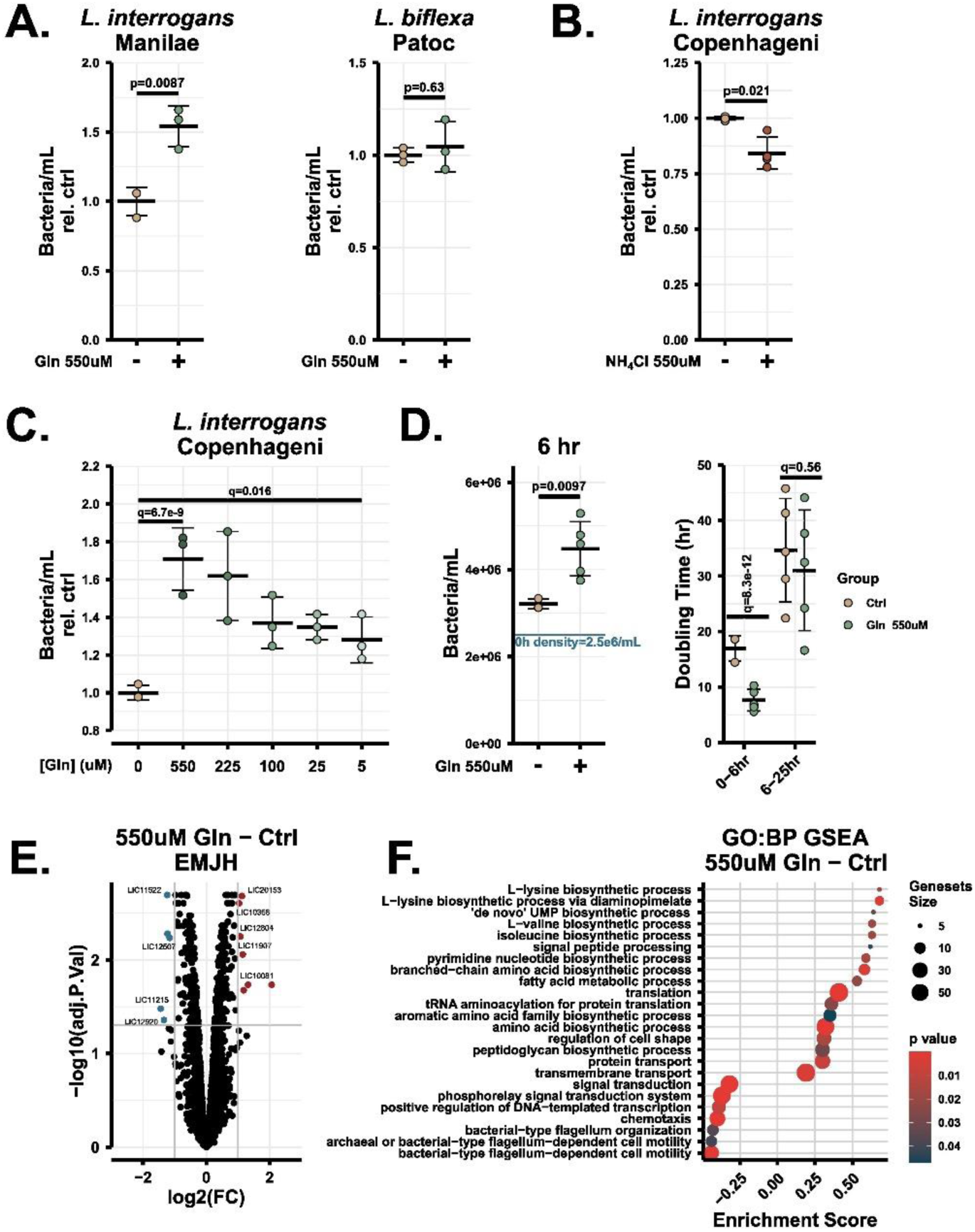
Exogenous glutamine stimulates a short-term proliferative boost in pathogenic *L. interrogans.* **A.** EMJH cultures of *L. interrogans* serovar Manilae (pathogenic) and *L. biflexa* serovar Patoc (nonpathogenic) supplemented with 550 µM Gln after seeding at 1e6 bacteria/mL, with proliferation assessed after one day by darkfield microscopy. **B.** Relative density after two days of proliferation from 2.5e6 bacteria/mL in EMJH cultures of *L. interrogans* serovar Copenhageni supplemented with an additional 550 µM ammonium chloride. **C.** Relative density after two days of proliferation from 2.5e6 bacteria/mL of EMJH cultures given the indicated amounts of Gln. **D.** (left) Relative density measurement six hours after seeding *L. interrogans* serovar Copenhageni at 2.5e6 bacteria/mL in EMJH supplemented with either vehicle or 550 µM Gln, demonstrating the early timescale of the effect. (right) From the same culture, doubling time between the 0-6 hour interval and the 6-25 hour interval. **E.** RNA-sequencing differential expression. RNA from *L. interrogans* exponential phase (5e7 bacteria/mL) EMJH cultures after four hours in EMJH+550 µM Gln versus EMJH (Gln-Ctrl). Red indicates increased transcript abundance, blue indicates decreased. **F.** Gene set enrichment analysis of expression trends in the Gln-Ctrl comparison. n=3 independent samples for A-C; n=5 for D; n=4 for Ctrl and n=3 for Gln in E-F. Welch’s t-test performed for A-B and D (left); for C and D (right) multiple contrasts, see Statistics in Methods. EMJH, Ellinghausen–McCullough–Johnson–Harris medium; Gln, glutamine; GO:BP, gene ontology: biological process; GSEA, gene set enrichment analysis; sHPLM, supplemented Human Plasma-Like Medium.

We also ascertained that the effect of Gln on proliferation was a medium-dependent phenomenon, as providing additional Gln atop the 550 µM already present in sHLPM did not accelerate the proliferation rate of *L. interrogans* serovar Copenhageni strain Fiocruz L1-130 (**S5A Fig**). Nitrogen availability is different between the two media, and since we identified that Gln served a metabolic role in sHPLM cultures, we next tested the possibility that the effect of Gln on proliferation in EMJH cultures was due to extra nitrogen supply. However, supplementing 550 µM ammonium to EMJH cultures did not phenocopy Gln (**Fig 4B**). This evidence suggested that the proliferative effect we observed was separate from the metabolic utilization of Gln-derived nitrogen to build biomass.

We suspected that Gln might be acting as a metabolite signal to accelerate proliferation in addition to serving a metabolic role as an important nutrient. While nutrients are required at high concentrations to support rapid consumption, signaling molecules can elicit effects at low concentrations. Consistent with this hypothesis, the addition of Gln at concentrations as low as 5 µM stimulated a disproportionately high increase in proliferation (∼1.25-fold above control) (**Fig 4C**). The Gln effect also appeared to be rapid and short-lived, as a decrease in doubling time manifested within six hours of Gln exposure, and thereafter the difference was muted (**Fig 4D** and **S5B Fig**). Metabolomics analysis revealed that the amount of Gln depleted within one day was minimal (**S5C Fig**), and thus consumption of the available Gln could not explain the transience of the effect. The low eliciting dosage, timescale, transience, and lack of correspondence to Gln levels in the medium support the model of a signaling metabolite.

To better understand its role as a potential signaling molecule, we assessed other aspects of *L. interrogans* physiology impacted by Gln. We examined biofilm formation and observed evidence that 550 µM Gln increased biofilm formation at 7 days and 14 days, whereas equimolar ammonium chloride failed to phenocopy this result, similar to in proliferation assays (**S5D Fig**). We also performed RNA-sequencing on *L. interrogans* EMJH cultures four hours after exposure to 550 µM Gln. Strikingly, we observed few genes meeting our fold change thresholds comparing Gln-exposed *L. interrogans* to vehicle controls (**Fig 4E** and **S3 Table**). We instead turned to gene set enrichment analysis and observed the upregulation of genes associated with various biosynthetic metabolic pathways, including some leading to cell wall synthesis, and translation. This matched well with the increased proliferation phenotype. There was also a general downregulation of signaling, motility, and chemotaxis-related genes (**Fig 4F**). Using genes with a gene ontology annotation or annotated in NCBI with a protein product related to transcriptional regulation, we identified possible regulatory proteins mediating the effect of Gln, according to the logic that many transcription regulation mechanisms are also self-regulating (**S5E Fig**). Overall, these results illustrate that environmental Gln can inform components of *L. interrogans* physiology involved in the pathogen’s lifecycle.

## Discussion

In this study, we developed two resources to advance metabolism research in *Leptospira* spp., both of which can be translated to the study of many microorganisms. First, we established a growth medium referred to as sHPLM, which approximates the human environment inhabited by infectious agents with a bloodstream phase. This medium induced phenotypes in *L. interrogans* similar to those observed *in vivo*. Second, we designed a workflow to perform metabolomics and stable isotope tracing on *L. interrogans* cultures and spent medium that enabled us to identify active metabolic pathways and qualitatively estimate the rates of nutrient transformation into biomass. The application of these resources revealed that Gln is a key source of nitrogen for *L. interrogans* under host-like conditions, where it is used to a similar extent as ammonium. Further investigation revealed that nitrogen assimilation can be chemically inhibited to block bacterial proliferation, and that Gln also has effects similar to an environmental signal, inducing proliferation, biofilm formation, and biosynthetic gene expression.

The disparity in phenotypes of *L. interrogans* cultured in EMJH and sHPLM emphasizes the need to shift culture techniques in microbiology to be more physiologically relevant and account for the strong influence of the environment on metabolic activity^40,41^. Indeed, there is an increasing call to use physiological medium for the study of infectious disease^41–44^. This trend has seen success in research on mammalian cells, where using HPLM has revealed key aspects of cancer therapy^24^ and immune cell activation^45^. The physiological accuracy of measurements made with *L. interrogans* cultured in this formulation was critical to our conclusions about host-relevant metabolism. We highlight that sensing of the environmental cues in sHPLM was sufficient to induce many virulence processes and metabolic alterations independent of physical interaction with host cells. As such, sHPLM can help deconvolute complex host-adaptation mechanisms with greater ease.

The complex formulation of sHPLM, which offered many potential nutrients, permitted a more representative assessment of the nutrient preferences of *L. interrogans* in hosts than could be achieved in minimal media such as EMJH, where only the bacterium’s capacity to adapt under duress can be evaluated^36,46^. This distinction is key to understanding what metabolites *L. interrogans* actually obtains from its host during infection, which may influence pathogenesis. From our list of putative nutrients, we focused on Gln due to its high concentration and its central role in nitrogen metabolism and host physiology^47^. Beyond Gln, our screen revealed several other metabolic characteristics of *L. interrogans* that were not pursued here. In line with conventional thinking, our results suggest that *L. interrogans* relies upon fatty acids. Thus, future work should examine which specific lipids are preferred under physiological conditions and the correspondence of this parasitic metabolism to the appearance of lipid-filled foamy macrophage phenotypes^48^ and injury-induced kidney steatosis^49,50^. Our screen also demonstrated Asp and pyruvate depletion by *L. interrogans*. Asp-derived nitrogen contributed minimally to *L. interrogans* biosynthesis, which additionally is evidence against its utilization as a carbon source^46^ and leaves its true role unknown. Prior work demonstrated that pyruvate neutralizes reactive oxygen species and other toxic oxidative products^51^, including those produced as byproducts of bacterial metabolism in *Escherichia coli*^52^. As *L. interrogans* relies on oxidative metabolism, future studies could assess whether this mechanism supports bacterial replication in the host. These connections to existing literature reinforce that our metabolomics-based nutrient screen can yield additional insights as a resource to the research community.

The metabolomics experiments performed herein represent, to our knowledge, the most thorough molecular interrogation of *L. interrogans* metabolism to date. Unlike many prior studies that have made inferences about *L. interrogans* metabolism from indirect measurements, we assess metabolism within bacterial cells directly by measuring the substrates and products of enzymatic reactions via LC/MS. Our method paves the way for more comprehensive multi-omics study designs in *Leptospira* cultures to better understand pathogenicity. Two other studies have used LC/MS in the context of leptospirosis to profile infected animal urine and plasma, but these primarily yielded information about the host response to infection^53,54^. It is important to note that our approach for sample isolation with vacuum filtration builds off of work in other liquid-cultured^55,56^ and solid-cultured species^57^. We previously employed a similar protocol to study gut microbe-host metabolic interaction^58^, but here the washing step is adjusted to effectively separate out metabolites from the complex formulation of sHPLM. This alteration is of critical importance when performing metabolomics on bacteria cultured in any nutrient-rich medium.

With this metabolomics method, we revisited the longstanding conclusion from *L. interrogans* Pomona that Gln could not serve as a sufficient nitrogen source unless first autoclaved to release ammonium^15^. Here, using *L. interrogans* Copenhageni, we demonstrated successful culture in medium lacking ammonium but containing unautoclaved Gln. By using ^15^N-tracers, we confirmed that Gln-derived nitrogen is used to produce biomass for proliferation and metabolite secretion. Metabolites secreted by *Leptospira* could contribute to biofilms, which are known to contain a substantial amount of extracellular DNA^59^. Alternately, this phenotype could be indicative of phage release. Our results necessitate a reconsideration of the published conclusion^60,61^ that ammonium is the only effective nitrogen source. Host innate immune cells, including macrophages, also rely upon Gln^62^, so our findings could imply metabolic competition in infected tissues. Our use of JHU-083 to target Gln metabolism, similar to past uses in *Mycobacterium tuberculosis* infection to limit bacterial burden and enhance macrophage responses^63^, showed promising inhibition of *L. interrogans* proliferation. This provides motivation for future preclinical studies using JHU-083 or the similar DON prodrug Sirpiglenastat that is already in phase 2 clinical trials (NCT07249372).

Based on our data, Gln also has a non-nutrient role in the *L. interrogans* lifecycle. Its rapid effect on proliferation and the transcriptome are consistent with a Gln-sensing axis. The abundance of Gln relative to ammonium varies between stages of the known lifecycle of pathogenic *Leptospira* spp. For example, Gln is high and ammonium is low in blood due to ammonia toxicity^64^, whereas the reverse is true in urine and possibly in the urine-contaminated soil and water that transmits *L. interrogans*^65^. It is noteworthy that the temporary proliferative bloom was observed in two pathogenic strains but not a nonpathogenic *Leptospira* strain. If this phenotype holds for other *Leptospira* spp., this could indicate that Gln metabolism and this signaling axis may have evolved to engage context-appropriate physiological responses in host-infecting *Leptospira* spp., similar to the virulence-activating effect of Gln on *Listeria monocytogenes*^66^. Further work should deconvolute the role of Gln from other host metabolites and resolve its effects in the timeline of host adaptation.

Collectively, using an innovative physiological medium and isotope tracing workflow to derive critical metabolic information, our data point to Gln as a key metabolite fueling *L. interrogans* replication and informing pathogen physiology. We hope that the present work will inspire the development of new therapeutic strategies for severe leptospirosis targeted at vulnerabilities in nitrogen metabolism.

## Methods

### Materials and handling of bacterial strains

*L. interrogans* serovar Copenhageni strain Fiocruz L1-130, *L. biflexa* serovar Patoc strain Patoc-1, and *L. interrogans* serovar Manilae strain L495 were generously provided by Dr. Jenifer Coburn, Medical College of Wisconsin. Pathogenic strains were at three passages from an animal host upon transfer and were used to no more than three additional passages to maintain virulence. Pathogenic strains were thawed rapidly in a 37 °C water bath in Hornsby-Alt-Nally (HAN) medium, prepared as previously described^67^, containing 100 µg/mL 5-fluorouracil (MilliporeSigma, F6627). Nonpathogenic *Leptospira* were thawed in liquid EMJH medium composed of 90% Base (BD Difco, 279410) suspended in Milli-Q water and autoclaved mixed with 10% Enrichment (BD Difco, 279510). Strains were frozen slowly in aliquots of 0.25 mL culture medium in a CoolCell chamber (Corning, 432001). Pathogenic strains were cultured either in liquid EMJH at 30 °C without shaking in conical tubes ∼75% filled with medium or at 37 °C and 5% CO2 in sHPLM or HAN in the same format, as required for successful culture in the respective media, and containing 100 µg/mL 5-fluorouracil. sHPLM culture medium was prepared by adding 10% 0.22 µm-filtered FBS (ThermoFisher, 16000044 or Corning 35-015-CV) and 2% BSA:oleate conjugate (Cayman Chemical, 29557) to HPLM (Life Technologies, A4899101). All bacterial strains were handled in a biosafety cabinet with BSL-2 procedures, respiratory protection, and restricted access. The metabolic inhibitors tested included dimethylammonium chloride (Fisher Scientific, AC155270050), methionine sulfoximine (MilliporeSigma, M5379), and JHU-083 (Cayman Chemical, 40642). Compounds supplemented to culture medium included ammonium chloride (MilliporeSigma, 213330), ^15^N-ammonium chloride (Cambridge Isotope Laboratories, NLM-467), L-glutamine (MilliporeSigma, 49419), ^15^N_2_-L-glutamine (Cambridge Isotope Laboratories, NLM-1328), urea (MilliporeSigma, U5128), ^15^N_2_-urea (Cambridge Isotope Laboratories, NLM-233), L-aspartic acid (MilliporeSigma, A8949), ^15^N-L-Aspartic acid (Cambridge Isotope Laboratories, NLM-718), and L-glutamic acid (MilliporeSigma, G8415).

### Isotope tracing

*L. interrogans* cultures, passaged for at least two days in sHPLM, were counted and centrifuged at 3700 RCF and 30 °C for 30 min, and the pellets were resuspended in 0.25 mL sHPLM per 40 mL of culture pelleted. During this time, fresh sHPLM was prepared and warmed to 37 °C, and 14 mL was removed for uninoculated replicates for each timepoint. The bacterial suspension was then recounted and inoculated into sHPLM at 1e8 bacteria/mL for 2-hour isolation, at 2.5e7 bacteria/mL for 24-hour isolation, and at 1e7/mL for 48-hour isolation and split into 15 mL conical tubes. The suspensions were rested for one hour in the incubator to allow metabolism to equilibrate, before each tube was sequentially spiked 1:1000 at ten-minute intervals with either ^15^N-labeled tracer or its ^14^N-unlabeled equivalent (uninoculated replicates were spiked with labeled tracer) and gently mixed by serological pipette twice. After the labeling time elapsed, a selection of inoculated tubes were counted to ensure a density around 1e8 bacteria/mL, and intracellular and medium samples were taken from each tube sequentially.

### Metabolite extraction from medium

After gentle mixing, 0.35 mL of culture was removed using a syringe, then filtered through a 0.1 µm filter (Millipore, SLVVR33RS), which sterilized media down to the limit of detection by darkfield microscopy and a Petroff-Hausser chamber. Filtered media was then diluted 1/10 in LC/MS-grade 2:2:1 acetonitrile:methanol:water pre-chilled at −25 °C, vortexed one min, incubated at −25 °C for one hour, snap-frozen in liquid nitrogen, and stored at −80 °C. Before analysis, samples were thawed on ice, centrifuged at 18000 RCF and 4 °C for ten min, and the supernatant transferred to glass LC vials.

### Metabolite extraction from cells

A vacuum filtration setup was prepared with a 500 mL Erlenmeyer filtration flask connected to a vacuum pump (KNF, UN811KTP) pulling −200 mbar. Capping the flask was the PELL magnetic filter funnel (Cytiva, 4247) with pre-activated (autoclaved milli-Q water) 47 mm 0.1 µm PTFE filter (Millipore, JVWP04700). *L. interrogans* cultures were poured over filters under vacuum, and afterwards 10 mL 0.85% saline (pre-warmed to 37 °C) dispensed around the sides of the reservoir to ensure movement of bacteria to the filter area. The filter was then removed and transferred to the bottom half of the Restek Diskcover-47 (VWR, R24020) which replaced the PELL filter holder unit on the vacuum trap. This next unit has a larger effective filtration area, upon which the filter was washed with 7 mL saline and gently blotted dry on the underside with a VWR Light-Duty tissue wipe. The filter was then transferred to a 50 mL conical tube containing 1.2 mL LC/MS-grade 2:2:1 acetonitrile:methanol:water, which had been pre-chilled on dry ice, then carefully vortexed for 30 sec without folding the filter, snap-frozen in liquid nitrogen for ten min (during which time the next sample was isolated) and transferred to dry ice until all samples were isolated. To process, conical tubes were sequentially thawed and sonicated 3 min at 20 °C, vortexed and shaken for 30 sec, snap-frozen in liquid nitrogen for three min, and re-sonicated three min at 20 °C, after which the liquid was removed to 1.5 mL tubes. These tubes were then vortexed briefly, snap frozen in liquid nitrogen for 3 min, then sonicated for 3 minutes and incubated at −25 °C overnight. It was confirmed that this procedure killed all bacteria. The following day, samples were centrifuged at 18000 RCF and 4 °C for 15 min and approximately 92.5% of the liquid transferred to fresh 1.5 mL tubes for lyophilizing on a SpeedVac concentrator. Dried pellets were resuspended in 60 µL 2:1 acetonitrile:water, subjected to two cycles of five min sonication and one min vortexing, then incubated at 4 °C overnight. Finally, samples were centrifuged at 18000 RCF and 4 °C for ten min and supernatant transferred to glass LC vials.

### Liquid chromatography mass spectrometry

For data collection, metabolite extracts were run on either a Thermo Q-Exactive Orbitrap or an ID-X II Orbitrap Mass Spectrometer interfaced with a Thermo Vanquish Horizon LC system using an iHILIC-(P) Classic 2.1 mm × 100 mm, 5 μm column (HILICON, 160.102.0520) with an iHILIC-(P) Classic Guard column (HILICON, 160.122.0520) attached. The column temperature was set to 45 °C. The mobile phases were as follows: for solution A, 95% water, 5% acetonitrile, 20 mM ammonium bicarbonate, 0.1% ammonium hydroxide solution (25% in water), 5 µM medronic acid; for solution B: 95% acetonitrile, 5% water. Each sample was injected (6 µL of intracellular and 2 µL of medium metabolite extract) and eluted with a linear gradient: at 250 µL/min 0-1 min 90% B, 1-12 min 35% B, 12-12.5 min 25% B, 12.5–14.5 min 25% B, 14.5–15 min 90% B, 15-20min at 400 µL/min and 90% B, and 20-22 min at 250 µL/min and 90% B for re-equilibration. MS/MS fragmentation data were collected using targeted methods at 20 eV. Metabolites were identified based on matching accurate mass, retention time, and MS/MS data to authentic reference standards in our in-house library or from curated databases for level one to two identification confidence. Chromatograms for selected metabolites were manually integrated in Skyline Daily (version 22.2.1.425).

### Isotope tracing analysis

Peak areas from manual integration in Skyline Daily (version 22.2.1.425) were loaded into R (version 4.4.2). Natural isotope abundance correction was performed in R using the github release of the accucore package^68^ (version 0.3.1), with a modified nitrogen_isotope_correction function to eliminate unnecessary nontracer corrections when complete separation of the labeled nitrogen isotopologue by the high-resolution mass spectrometer was obtained (confirmed in Skyline Daily). Isotopologue peak areas were normalized as a percent of the summed intensity of all isotopologue peaks per sample and metabolite, and plotted as 1) a summed total percentage of all labeled (M>0) isotopologues, 2) as the percentage of each isotopologue, or 3) as a percentage of total ^15^N nitrogen atoms in the metabolite pool with correction for the initial tracer labeling percentage to reflect the true percent of nitrogen atoms derived from the tracer.

### Depletion analysis from metabolomics data

Peak areas from manual integration in Skyline Daily (version 22.2.1.425) were loaded into R (version 4.4.2). First, metabolites were normalized as zero-centered fold change relative to the average signal from day one samples. Uninoculated control measurements were moderated to reduce measurement error by using linear models created from metabolite-specific peak area data and obtaining predicted outputs for days one, three, and five, which were then normalized as zero-centered fold change relative to the predicted value from the day one sample. Fold changes in inoculated samples were then corrected for changes in the uninoculated controls by subtracting day-matched uninoculated values from inoculated sample values. Fold changes from days one through five were plotted as a histogram to choose a suitable cutoff, and metabolites meeting this cutoff with a q-value<0.05 as determined by Welch’s t-test with Benjamini-Hochberg adjustment were plotted in R. Metabolomics data have been deposited in the NIH Metabolomics Workbench repository as of the time of publication.

### Density measurements and proliferation assay

Bacterial density was enumerated with a Petroff-Hausser chamber (Electron Microscopy Sciences, 63512-24) loaded with 2.5 µL culture, with dilution in HAN, EMJH or DI water as needed, visualized in darkfield on a Leica DM500 microscope (NCI Inc., 13613215). Between four and eight grid sections of the 1 mm center square were counted at 400X total magnification, averaged, and multiplied by 25 grid squares by 50000 mL^-1^ to obtain the bacterial culture concentration (bacteria/mL). For proliferation assays, bacteria were resuspended to the indicated density in fresh medium and aliquoted at 3-4 mL each to 5 mL tubes (Midsci, C2535). Tube caps were parafilm-wrapped and incubated for the indicated times, whereupon culture was gently mixed to resuspend sedimented bacteria and counted.

### Biofilm assay and microscopy

*L. interrogans* culture was dispensed at 150 µL/well in 96-well plates (Midsci, 781962) with outer wells filled with deionized water and incubated for seven days or 14 days. Medium was carefully removed, and 175 µL 10% Formalin added and incubated 30 min at room temperature before removal. Each well was then washed twice with 200 µL PBS (Gibco, 10-010-049), and 175 µL Crystal Violet Solution added and incubated for ten min. Each well was then washed twice with 200 µL PBS, and the plate was dried overnight. Individual wells were imaged using a Leica THUNDER imaging system at 5x and 10x with automatic stitching together. Images were imported into ImageJ (version 1.50i), clipped to the boundaries of the well, color thresholding applied (HSB: Hue 39-255, Saturation 50-255, Brightness 99-199), and area measured relative to 1 mm scale bars.

### RNA isolation

*L. interrogans* cultures were centrifuged at 3700 RCF and 30 °C for 30 min, then the pellet resuspended in 1 mL Trizol (Invitrogen,15596018) and snap-frozen in liquid nitrogen for storage at −80 °C. For processing, samples were thawed on ice, centrifuged for five min at 18000 RCF and 4 °C, 0.2 mL chloroform added and inverted several times to mix before incubation at room temperature for three min. Samples were centrifuged for 15 min at 12000 RCF and 4 °C and the clear aqueous phase transferred to a new tube, to which was added 0.5 mL cold isopropanol. Tubes were inverted gently to mix and incubated for ten min on ice before being centrifuged for 15 min at 18000 RCF and 4 °C. The supernatant was removed and the RNA pellet resuspended in 75% ethanol, vortexed briefly, and centrifuged again for 10 min at 18000 RCF and 4 °C. The ethanol was then carefully removed, the pellet dried for five min at room temperature and resuspended in nuclease-free water by pipetting and heating at 55 °C for five min. RNA was then treated with DNase as per the instruction of the DNA-free kit (ThermoFisher, AM1906) and quantified with a NanoDrop instrument (ThermoFisher).

### RNA-sequencing and analysis

RNA-sequencing was performed by the Genome and Technology Access Center (GTAC) at Washington University in St. Louis. RNA integrity was quantified with the Agilent Bioanalyzer or 4200 Tapestation. cDNA libraries were prepared with an input of 500 ng to 1 µg of total RNA, as follows. RNA was fragmented according to the FastSelect protocol (Qiagen) in reverse transcriptase buffer (94 °C for five min, 75 °C for two min, 70 °C for two min, 65 °C for two min, 60 °C for two min, 55 °C for two min, 37 °C for five min, 25 °C for five min), with ribosomal RNA depleted during the process. cDNA was synthesized from remaining mRNA using the SuperScript III RT enzyme (Life Technologies) and random hexamers according to manufacturer’s instructions. ds-cDNA was formed using a second strand reaction. The product library was blunt ended and had an A base added to the 3’ ends before then ligating Illumina sequencing adapters. The library was amplified for 15 cycles using primers incorporating unique dual index tags. Sequencing was performed on an Illumina NovaSeq X Plus using paired end reads extending 150 bases, with Illumina’s bcl2fastq software used for base calling and demultiplexing setting a threshold of a maximum of one mismatch in the indexing read. Reads were aligned with STAR (version 2.7.11b) to the *Leptospira interrogans* serovar Copenhageni str. Fiocruz L1-130 GCF_000007685.1 (ASM768v1) primary assembly. Gene counts were calculated from uniquely aligned and unambiguous fragments using Subread:featureCount (version 2.0.8). Sequencing performance was evaluated based on total versus uniquely aligned reads, features detected, the ribosomal fraction, known junction saturation, and read distribution over known gene models using RSeQC (version 5.04).

Differential gene count was evaluated with the R/Bioconductor (version 4.4.0) package EdgeR and Limma using TMM normalization size factors to adjust for library size. Reads for ribosomal genes and those with a lower count than one count per million in at least three samples were excluded. A Limma generalized linear model was fitted using weighted likelihoods calculated from gene and sample-wise mean-variance relationships and the voomWithQualityWeights function. This model was used for differential expression analysis, with Benjamini-Hochberg false-discovery rate adjustment applied for p-values. One sample in the EMJH+Gln group was determined to be a statistical outlier and was removed from the analysis, leaving three independent replicates. The data are stored under NCBI BioProject number PRJNA1401465 and GEO accession GSE316384.

Gene set analyses were performed in R (version 4.4.2) using the packages UniProt.ws (version 2.46.1), ClusterProfiler (version 4.14.6) and functions from the package HypeR (version 2.0.0) modified to employ a two-tailed version of the ks.test function. Gene sets were downloaded from the QuickGO website^70^ with filters: taxon=173 (include descendants), gene product=proteins. GO terms were further filtered to exclude sets with fewer than five genes. The remaining library of sets contained many sets with few detected genes, which skewed post-hoc p-value correction; thus, p-values are reported for gene set analysis.

### RT-qPCR

RNA was first reverse transcribed to cDNA using the iSCRIPT cDNA Synthesis Kit (Bio-Rad, 1708891) according to the manufacturer’s protocol and a Bio-Rad S1000 Thermal Cycler (Bio-Rad, 1852148). cDNA was diluted by 1/4 using nuclease-free water. Primers were synthesized by IDT. Sequences: 16S-F 5’-AGGTAAAGATTCGATAGGAGCG-3’, 16S-R 5’-GTAGAACCGCTTTCTGGGG-3’, glnA-F 5’-CTCTTCTTGCCTTGTCTCCGT-3’, glnA-R 5’- GTGGGGTTCTTTCTCACGCT-3’. Primers were resuspended to 100 nM in nuclease-free water and diluted to working concentrations of 10 nM. qPCR reactions were assembled in 96-well reaction plates (Applied Biosystems, 4346906) as follows: 2 µL diluted cDNA, 0.5 µL each forward and reverse primer, 5 µL Power SYBR PCR Master Mix (Applied Biosystems, 4367659), and 2 µL nuclease-free water. Each plate was centrifuged briefly at 2000 RCF, then run on an Applied Biosystems StepOne Plus instrument (software version 2.3) in technical duplicate. Gene expression was calculated as ΔCt relative to the 16S RNA measurement for each sample.

### Statistical analysis

Statistical analyses were performed in R (version 4.4.2). Data were analyzed by two-tailed Welch’s T-test if only one pairwise contrast was made. If multiple pairwise contrasts were made, groupwise data were assessed for heteroscedasticity with a Bartlett test, cutoff p<0.1, and either a hetero- or homoscedastic generalized least squares model created using the gls function from the nlme package (version 3.1-166). Prespecified pairwise contrasts from the model were tested using the glht function from the multcomp package (version 1.4-26), analogous to specified contrasts tested on an anova model^69^. Post-hoc correction using the Benjamini-Hochberg method was then applied to produce q-values. Effect sizes were calculated with the effectsize package (version 1.0.0). A pairwise difference was considered statistically significant when p<0.05 or q<0.05 and effect size was ≥2.5. Data in all point plots are presented as mean +/- 1 standard deviation. All plots were created in R, using packages: ggplot2 (version 3.5.1), NatParksPallettes (version 0.2.0), ggh4x (version 0.2.8), plyr (version 1.8.9), dplyr (version 1.1.4), ComplexHeatmap (version 2.22.0), circlize (version 0.4.17), ggrepel (version 0.9.6), enrichplot (version 1.26.6), and reshape2 (version 1.4.4). Schematics were created using Inkscape or Biorender.

## Acknowledgements

We thank Dr. Jenifer Coburn for generously sharing bacterial strains and protocols. We thank the Genome Technology Access Center at the McDonnell Genome Institute at Washington University School of Medicine for help with transcriptomic analysis. The Center is partially supported by NCI Cancer Center Support Grant #P30 CA91842 to the Siteman Cancer Center from the National Center for Research Resources (NCRR), a component of the National Institutes of Health (NIH), and NIH Roadmap for Medical Research. This publication is solely the responsibility of the authors and does not necessarily represent the official view of NCRR or NIH.

## Author Contributions

MHW, LPS, and GJP designed the study. MHW and NS carried out experiments. MHW analyzed data, prepared manuscript figures, and wrote the manuscript. LPS and GJP provided resources, funding, and conceptual input for experiments and supervised the research. All authors reviewed, edited, and approved this manuscript.

---

**Supplementary Figure 1.**
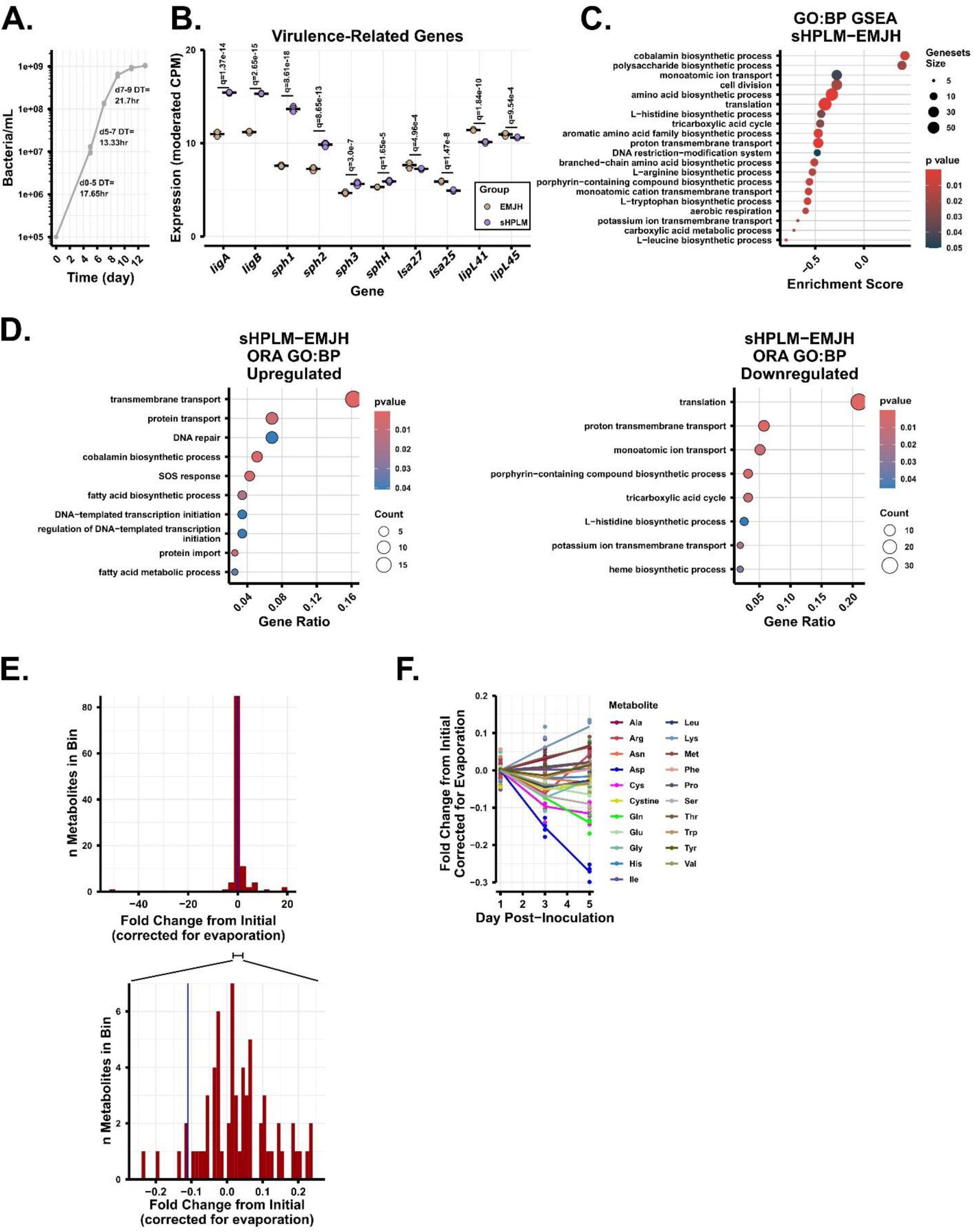
sHPLM cultures mimic the *in vivo* transcriptome and consume select metabolites. **A.** Growth curve of *L. interrogans* Copenhageni strain Fiocruz L1-130 grown at 30 °C in liquid EMJH from 1e5 bacteria/mL. **B.** CPM transcript expression in selected virulence-related genes from EMJH cultures switched to EMJH or sHPLM for four hours. **C.** GSEA of expression trends in the sHPLM-EMJH comparison. **D.** Gene set ORA in q<0.05 and log2(FC)>1 or log2(FC)<-1 genes in the sHPLM-EMJH comparison. **E.** (upper) Histogram of metabolite total fold changes from 111 detected metabolites from days one through five. (lower) Zoomed-in plot of histogram showing −11% threshold. **F.** Line plots of amino acid fold changes over time. n=4 independent samples per group for A-F. Statistical difference in B determined with EdgeR and Benjamini-Hochberg correction. Ala, alanine; Arg, arginine; Asn, asparagine; Asp, aspartic acid; CPM, counts-per-million; Cys, cysteine; DT, doubling time; EMJH, Ellinghausen–McCullough–Johnson–Harris medium; Gln, glutamine; Glu, glutamic acid; Gly, glycine; GO:BP, gene ontology: biological process; GSEA, gene set enrichment analysis; His, histidine; Ile, isoleucine; Leu, leucine; Lys, lysine; Met, methionine; ORA, overrepresentation analysis; Phe, phenylalanine; Pro, proline; Ser, serine; sHPLM, supplemented human plasma-like medium; Thr, threonine; Trp, tryptophan; Tyr, tyrosine; Val, valine.

**Supplementary Figure 2.**
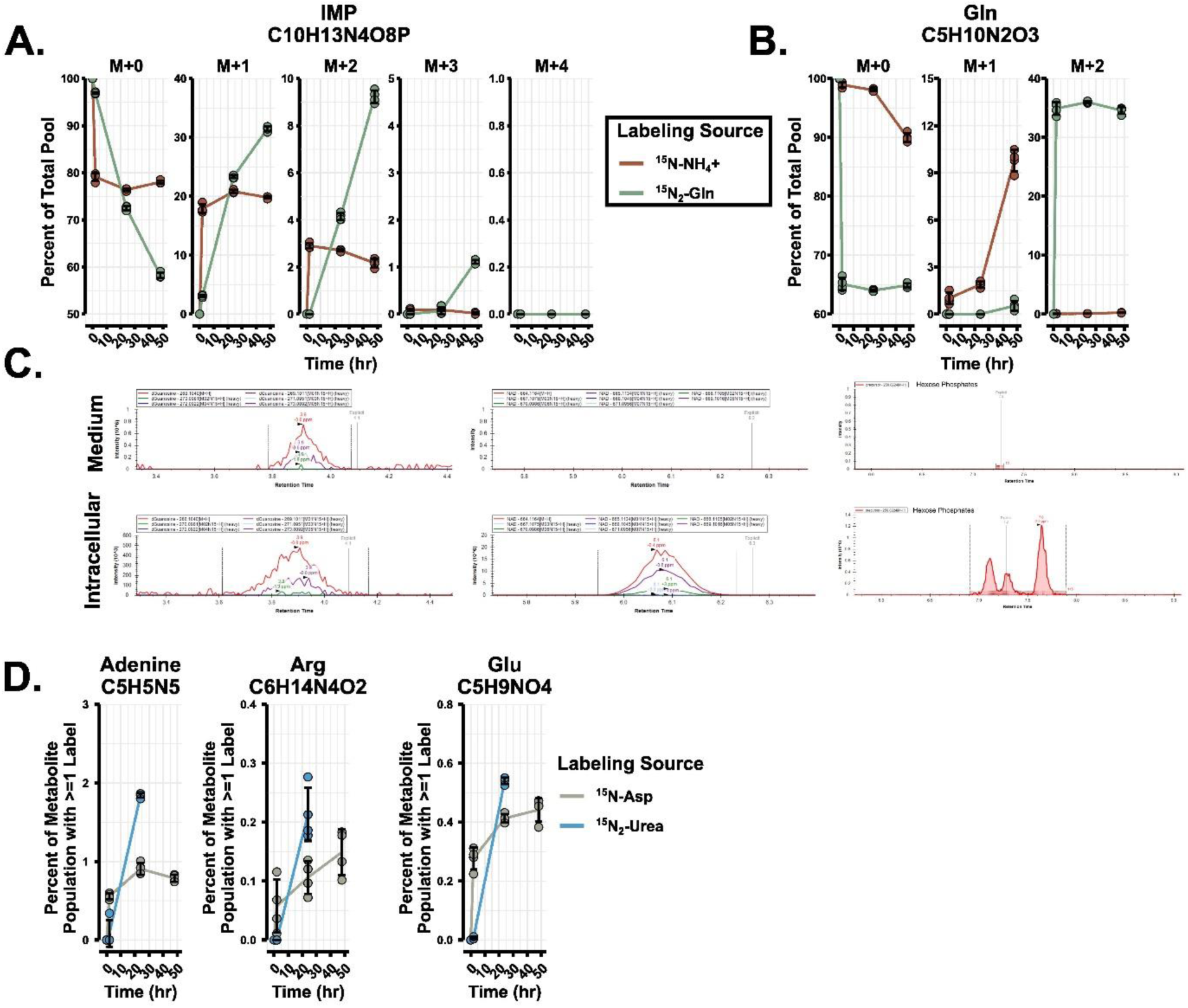
Stable isotope tracing of nitrogen sources in physiologically cultured *L. interrogans*. **A.** Example breakdown of relative abundance of isotopologues (mass isotopologue distribution) for IMP over time, showing gradual trickling of ^15^N into downstream metabolism. Data from experiments shown in Fig 2. **B.** Mass isotopologue distribution of Gln labeled from either exogenous ammonium or Gln. Note that some medium contamination may lead to overestimation of M+2 label from exogenous Gln and underestimation of M+1 and M+2 label from ammonium. **C.** Medium and intracellular metabolite extracted ion chromatograms for *m/z* corresponding to dGuanosine (left), NAD (middle), and hexose phosphate (right). Retention time (x-axis) in minutes. **D.** Percentage of given metabolite populations that have at least one labeled nitrogen from host-like medium-derived ^15^N-Asp or ^15^N_2_-urea after 2, 24, and 48 hours of labeling in sHPLM spiked to a concentration of 33.3% labeled nitrogen source. Metabolites are representative of initial assimilation (Glu), purine biosynthesis (adenine), and arginine biosynthesis (Arg), with the criteria that they must be detected in both experiments. Asp labeling data is from the experiment shown in Fig 2B-C. Data in A-B from experiments shown in Fig 2. n=4 independent samples per group in A, B, D. Asp, aspartic acid; dGuanosine, deoxyguanosine; Gln, glutamine; IMP, inosine monophosphate; NAD, nicotinamide adenine dinucleotide.

**Supplementary Figure 3.**
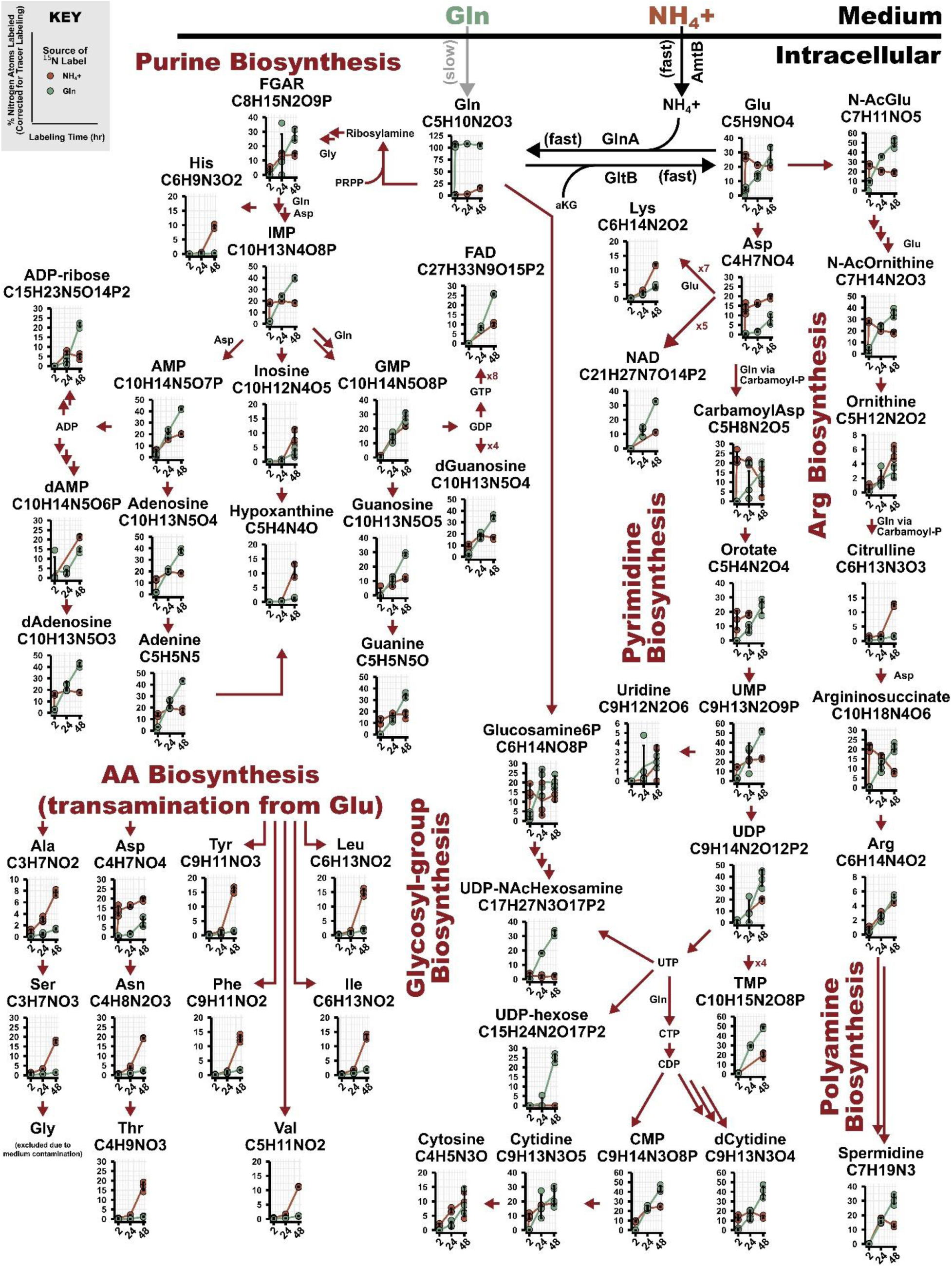
Metabolic map of nitrogen utilization in *L. interrogans.* Mapping of nitrogen flux through detected metabolites in sHPLM-cultured *L. interrogans*, showing active pathways and reactions. Based on isotope tracing data with ammonium chloride, Gln, Asp, and urea as precursors. Red arrows indicate reactions along pathways, corresponding to a single enzymatic step. Some metabolites were not detected at specific timepoints and were omitted. Percent labeling of nitrogen atoms calculated based on mass isotopomer distribution for each metabolite and divided by 1/3 to reflect the true percentage of nitrogen derived from each tracer. Zero-hour timepoint imputed at 0% labeling. n=4 independent samples per tracer per time-point. AA, amino acid; ADP, adenosine diphosphate; ADP-ribose, adenosine diphosphate ribose; aKG, alpha-ketoglutarate; Ala, alanine; AMP, adenosine monophosphate; Arg, arginine; Asn, asparagine; CarbamoylAsp, carbamoyl aspartate; Carbamoyl-P, carbamoyl phosphate; CDP, cytidine diphosphate; CMP, cytidine monophosphate; CTP, cytidine triphosphate; dAdenosine, deoxyadenosine; dAMP, deoxyadenosine monophosphate; dCytidine, deoxycytidine; dGuanosine, deoxyguanosine; FAD, flavin adenine dinucleotide; FGAR, 5’-phosphoribosyl-N-formylglycinamide; GDP, guanosine diphosphate; Gln, glutamine; Glu, glutamic acid; Glucosamine6P, glucosamine-6-phosphate; Gly, glycine; GMP, guanosine monophosphate; GTP, guanosine diphosphate; His, histidine; IMP, inosine monophosphate; Ile, isoleucine; Leu, leucine; Lys, lysine; NAD, nicotinamide adenine dinucleotide; N-AcGlu, N-acetylglutamic acid; N-AcOrnithine, N-acetylornithine; Phe, phenylalanine; PRPP, phosphoribosyl pyrophosphate; Ser, serine; Thr, threonine; TMP, thymidine monophosphate; Tyr, tyrosine; UDP, uridine diphosphate; UMP, uridine monophosphate; UDP-hexose, uridine diphosphate hexose; UDP-NAcHexosamine, uridine diphosphate N-acetylhexosamine; UTP, uridine triphosphate; Val, valine.

**Supplementary Figure 4.**
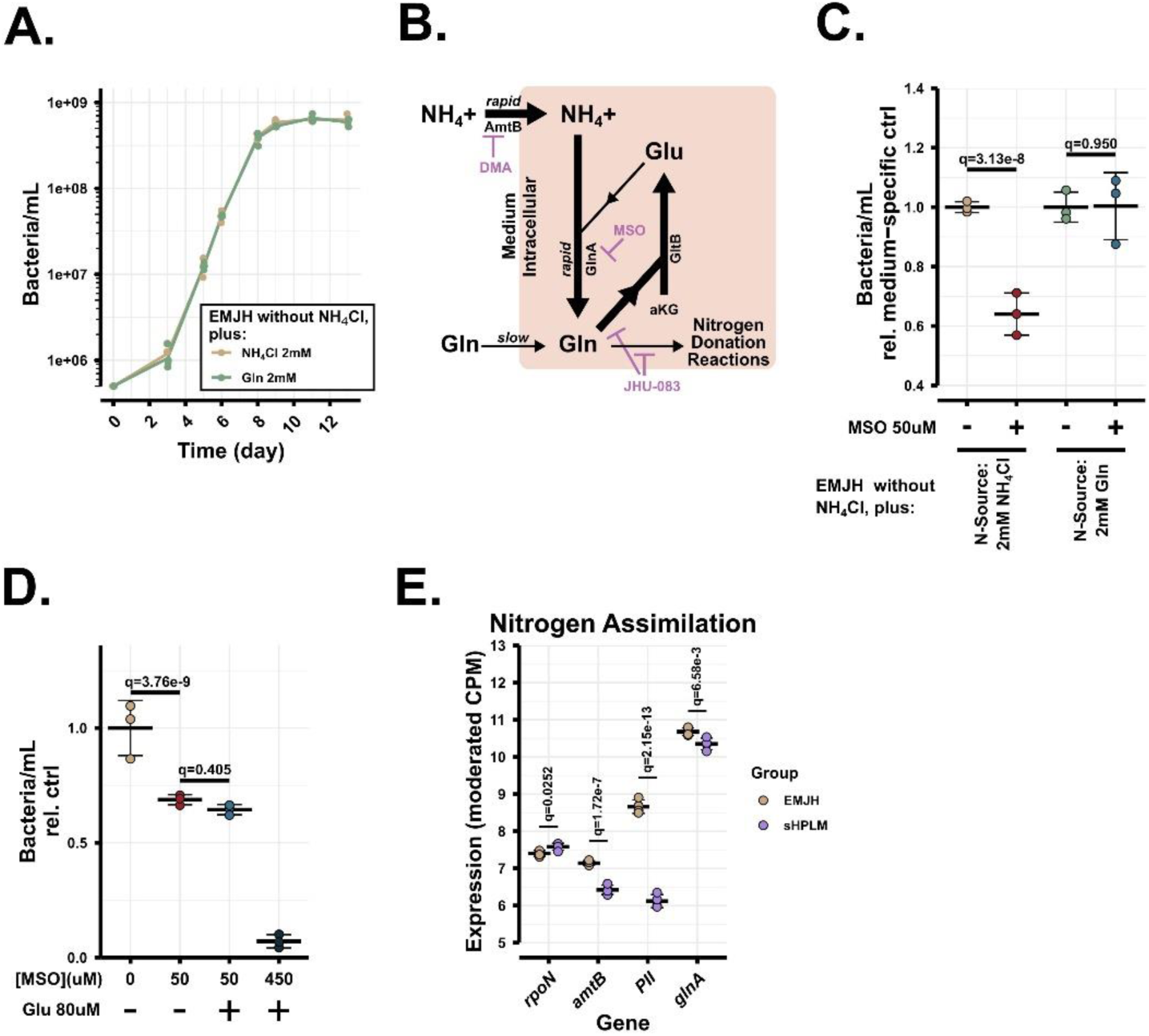
Chemical inhibition and medium effects on nitrogen metabolism. **A.** Growth curve of *L. interrogans* Copenhageni grown from 5e5 bacteria/mL in EMJH lacking ammonium chloride, complemented back with either 2 mM ammonium chloride or Gln. **B.** Points of chemical intervention in nitrogen metabolism. **C.** Two-day proliferation assay of *L. interrogans* grown on custom EMJH formulated without ammonium chloride and re-supplemented with either 2 mM ammonium chloride or Gln and treated with 50 µM MSO, a glutamine synthetase inhibitor. Density measured by darkfield microscopy two days after seeding at 2.5e6 bacteria/mL. **D.** Proliferation assay of EMJH cultures given 50 or 450 µM MSO and 80 µM L-glutamic acid, measured at four-day endpoint from inoculation at 2.5e6 bacteria/mL. **E.** CPM transcript expression in selected nitrogen assimilation-related genes from EMJH cultures switched to EMJH or sHPLM for four hours. Data from experiment presented in Fig 1B. n=4 independent samples per group for A; n=3 for B-D; n=4 for E. For C-E multiple contrasts, see Statistics in Methods. CPM, counts-per-million; aKG, alpha-ketoglutarate; DMA, Dimethylammonium; EMJH, Ellinghausen–McCullough–Johnson–Harris medium; Glu, glutamic acid; Gln, glutamine; MSO, methionine sulfoximine; NH_4_Cl, ammonium chloride; sHPLM, supplemented Human Plasma-Like Medium.

**Supplementary Figure 5.**
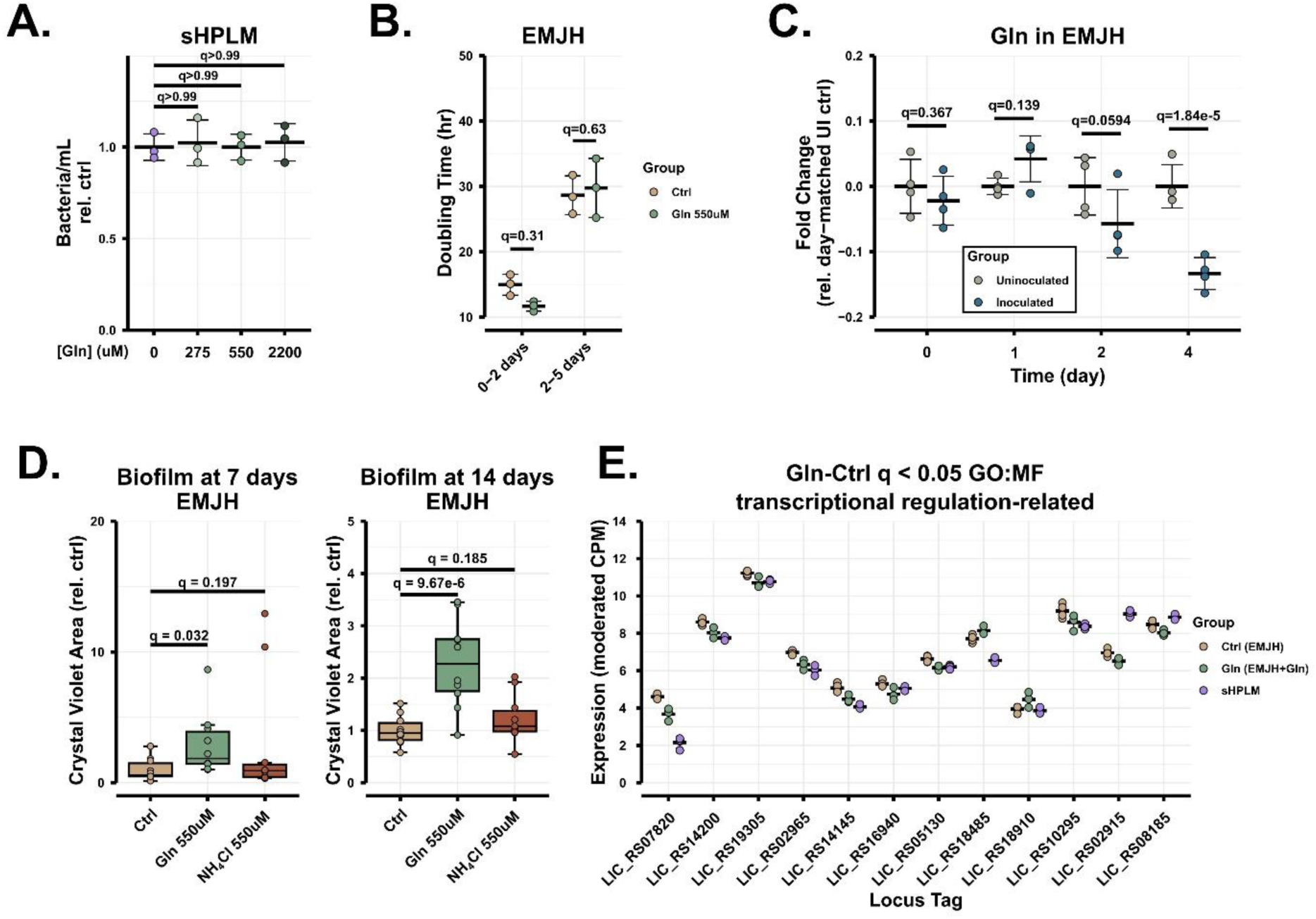
The effect of glutamine on *L. interrogans* physiology. **A.** Proliferation assay of sHPLM cultures given the indicated amounts of Gln (in addition to the 550 µM inherent in HPLM), with density measured by darkfield microscopy at day three from an inoculated density of 1e6 bacteria/mL. **B.** Calculated doubling time in experiment presented in Fig 3D. Counts taken at two and five days. **C.** LC/MS measurements of relative Gln concentration over time in EMJH cultures supplemented with 550 µM Gln. Normalized to average signal in day-matched UI evaporation controls. **D.** Crystal violet-stained biofilm quantitation at seven and 14 days after inoculation at 2.5e6/mL in 96-well plate with vehicle, 550 µM Gln, or 550 µM ammonium chloride. **E.** CPM transcript expression in *L. interrogans* EMJH cultures four hours after switching to fresh EMJH, EMJH+550 µM Gln, or sHPLM in q<0.05 genes with gene ontology annotations containing the keywords “sigma factor”, “DNA binding”, “DNA-binding”, “transcription factor”, “transcriptional regulator”, or “transcriptional repressor.” n=3 independent samples per group in B and Gln group of E, n=4 in C and Ctrl and sHPLM groups of E, n=10 in D. For A-D multiple contrasts, see Statistics in Methods. Statistical difference in E determined with EdgeR and Benjamini-Hochberg correction. CPM, counts per million; EMJH, Ellinghausen–McCullough–Johnson–Harris medium; GO:MF, gene ontology molecular function; NH_4_Cl, ammonium chloride; sHPLM, supplemented Human Plasma-Like Medium; UI, uninoculated.

